# Exploring the accuracy of ab initio prediction methods for viral pseudoknotted RNA structures

**DOI:** 10.1101/2024.03.21.586060

**Authors:** Vasco Medeiros, Jennifer M. Pearl, Mia Carboni, Ece Er, Stamatia Zafeiri

## Abstract

The prediction of tertiary RNA structures is significant to the field of medicine (e.g. mRNA vaccines, genome editing), and the exploration of viral transcripts. Though many RNA folding software exist, few studies have condensed their locus of attention solely to viral pseudoknotted RNA. These regulatory pseudoknots play a role in genome replication, gene expression, and protein synthesis. This study explores five RNA folding engines that compute either the minimum free energy (MFE) or the maximum expected accuracy (MEA). These folding engines were tested against 26 experimentally derived short pseudoknotted sequences (20-150nt) using metrics that are commonly applied to software prediction accuracy (e.g. F_1_ scoring, PPV). This paper reports higher accuracy RNA prediction engines, such as pKiss, when compared to previous iterations of the software, and when compared to older folding engines. They show that MEA folding software does not always outperform MFE folding software in prediction accuracy when assessed with metrics such as percent error, sensitivity, PPV, and F_1_ scoring when applied to viral pseudoknotted RNA. Moreover, the results suggest that thermodynamic model parameters will not ensure accuracy if auxiliary parameters such as Mg^2+^ binding, dangling end options, and H-type penalties are not applied. The observations reported in this paper highlight the quality between different *ab initio* prediction methods while enforcing the idea that a better understanding of intracellular thermodynamics is necessary for a more efficacious screening of RNAs.

**Importance:** The importance of accurately predicting RNA structures cannot be overstated, particularly in the context of viral biology and the development of therapeutic interventions such as mRNA vaccines and genome editing. Our study addresses the gap in the existing literature by concentrating solely on viral pseudoknotted RNA, which plays a crucial role in viral replication, gene expression, and protein synthesis. Our study sheds light on the debate surrounding minimum free energy (MFE) versus maximum expected accuracy (MEA) models in RNA folding predictions. Contrary to existing beliefs, we found that MEA models do not consistently outperform MFE models, especially in the context of viral pseudoknotted RNAs. Our research contributes to advancing the field of computational biology by providing insights into the efficacy of different prediction methods and emphasizing the need for a deeper understanding of intracellular thermodynamics to improve RNA structure predictions.

## Introduction

Computational biology is one of the key tools we possess to understand RNA folding, and is employed in pharmacokinetics, drug discovery, and pharmacology. In silico predictions of catalytic RNAs help narrow down and consolidate a surfeit of data while expediting the search for potential drug targets. As of now, we know that catalytic RNA controls for ribozymes, riboswitches, mRNA vaccines, thermosensors, and essential elements of genome editing [1–4]. This is due to RNA’s ability to fold itself into tertiary structures (pseudoknots), forming binding pockets, and active site clefts which can act as targets for active pharmaceutical ingredients (APIs) [5]. As AI, computer processing, and data thruput continue to advance, we are witnessing these methodologies more frequently implemented in the fields of virology, and viral RNA [6]. This paper will explore the different ways in which stochastic folding engines predict viral pseudoknotted RNAs and the accuracy of these approaches.

Pseudoknots are structural motifs found in almost all classes of RNA. While most RNA forms planer secondary structures, these three-dimensional structures embody up to 30% of tertiary non-planer motifs in G + C-rich RNA sequences [7]. In the context of viruses, these pseudoknots control gene expression and protein synthesis. Many catalytic RNAs regulate the mechanisms of action associated with viral replication, viral translation, or both. An example of this can be seen in satellite viruses, (e.g. hepatitis delta virus, satellite tobacco necrosis virus 1) which encode ribozymes that are folded by pseudoknotted structures [8,9].

For decades, scientists have explored and created different prediction software to better elucidate the complicated nature of RNA folding. Though RNA is a biopolymer that folds in a specific manner, it can be difficult to discern which secondary prediction algorithms, alignment sequences, or applied mathematics would result in the most accurate model. This difficulty becomes more apparent when noting how much the field has changed over the years, and how small changes in the underlying formalisms and constraints can result in drastic differences in the final predicted structure.

RNA folding occurs through populated intermediates and is accomplished in a hierarchical manner where secondary planer forms come prior to tertiary contacts [10]. This allows software engines to model both canonical and non-canonical base pairs, making it so the inputs within V(i,j) base pairs have a range of integer values (rather than binary values) dependent on the base pairs they form [11,12]. Using mathematical formalisms, derived from prior experimental data to model the effects of salinity, pH, temperature, loop entropies, and stacking formations, we can generate a “Pseudo-Energy Model’. This grants us a measure of the relative probability of different RNA secondary structures, expounding on the ensemble free energy, and the equilibrium concentrations of all possible structures, all of which correspond to the topological character of the RNA strand [13].

Within the RNA template, the first base at the 3’ terminus is regarded as 1, and the final base found at the 5’ terminus is regarded as N. In the total secondary structure, made up of V(i,j) base pairs, the index 1 ≤ i < j ≤ N should be set. Each integer within a given matrix will represent the ith nucleotide being paired with the jth nucleotide. The base pairs, (G-C), (A-U), and at times, (G-U) (dependent on the algorithm/software used) are the integer values that contribute to the matrix field. The entire structure of length N is regularly represented in Feynman diagrams, also known as arc and chord diagrams (**Fig. 1A**), where each nucleotide is represented as a point on the chain, while each arc represents a base pair forming between any nucleotides i and j.

**Figure 1.**
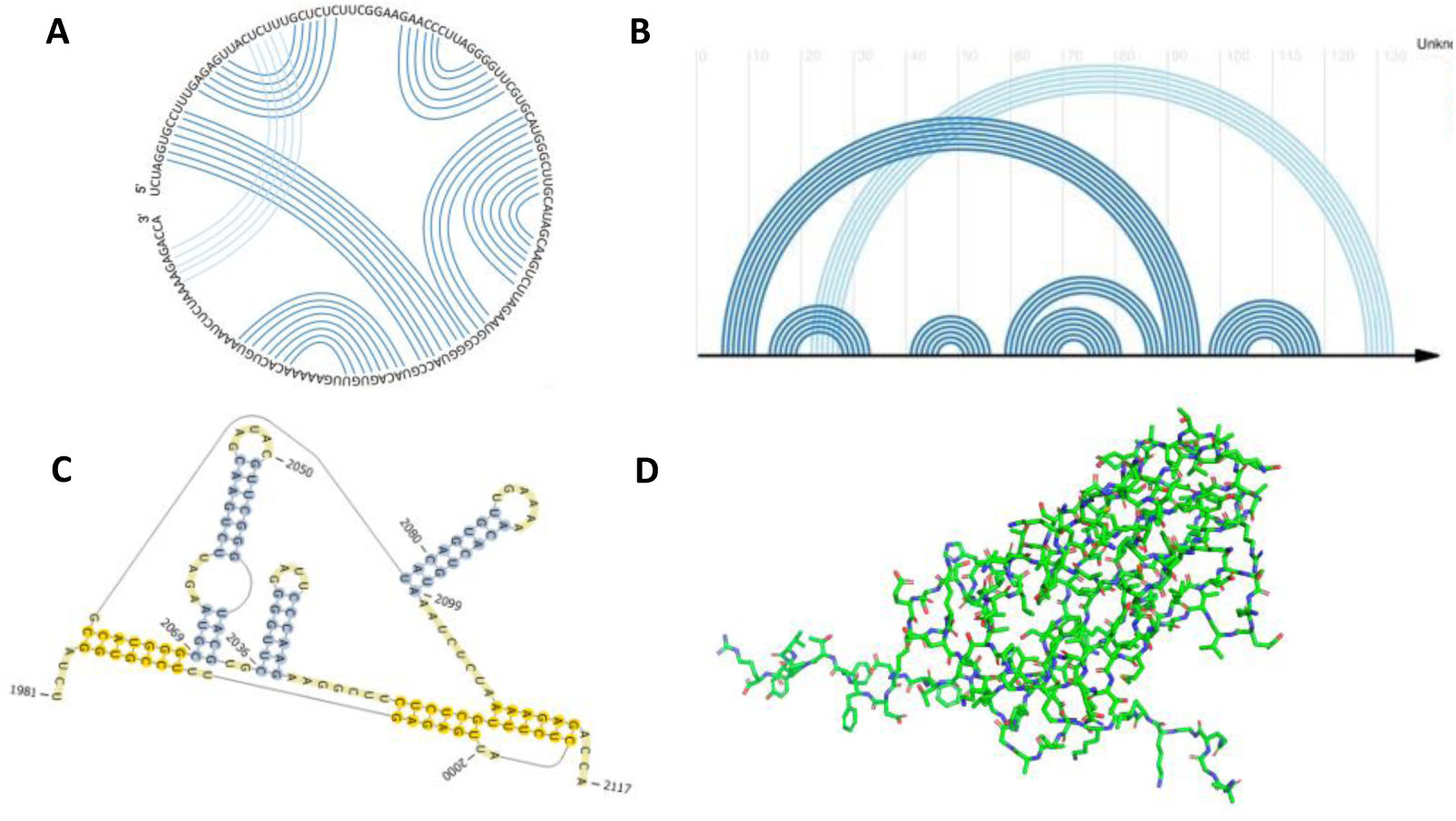
Ways in which to model pseudoknotted RNA. (**A**) Circular arc and cord diagram of the viral tRNA-like brome mosaic virus [14,15]. (**B**) Planer arc and chord drawing of the viral tRNA-like brome mosaic virus made with R-chie package [16]. (**C**) Planer representation of viral tRNA-like brome mosaic virus using nitrogenous bases [17]. (**D**) 3D model of tRNA-like brome mosaic virus in stick format made using PyMOL Molecular Graphics System, Version 2.0 [18].

One definition of “a pseudoknot” could be: “a template of RNA in which nucleotides within a loop pair with regions that do not pertain to the helices that close said loop.” Another definition could be “an RNA secondary structure that forms base pair regions upstream or downstream, resulting in stem-loop structures.” Their topology is more varied than most other assemblies of RNA, presenting a challenge for in silico prediction software.

While comparative approaches exist in the solving of optimal RNA structures [19,20], including web servers such as KNetFold [21], and pAliKiss [22], this review will focus on the *ab initio* method-topological predictions based primarily on the RNA secondary structure. The accuracy of these *ab initio* stochastic RNA folding software will be assessed in relation to a catalog of 26 distinct viral pseudoknotted RNAs taken from PseudoBase++ [23], whose wild-type structures have been previously determined via sequence com-parison, structure probing, mutagenesis, and NMR. The RNA predictions generated in this paper, imparted by the underlying formalisms of the software, will result in structures that represent either the minimum free energy (MFE) or the maximum expected accuracy (MEA), as both models are compared.

It can generally be posited that base pairs, when formed, lower the Gibbs free energy of a ribonucleic strand, making use of the attractive interactions between the complementary strands. MFE prediction algorithms assess these by solving for the maximum number of nucleotide pairings, via their thermodynamic properties, which generally results in the lowest energy form.

It is important to note however that, in nature, kinetic barriers, environmental conditions, and other factors may influence RNA folding intermediates, resulting in a physiologically favored RNA that does not coincide with the minimum free energy MFE structure [24]. Conversely, MEA models compute the final RNA structure via a partition function (a function that is used to calculate the thermodynamic properties of a system) that implements hard and soft constraints based on electrostatic interactions, stacking interactions, adjacent complementary base pairs, and other variables depending on the software in question [24]. This results in a final structure that may not necessarily encompass the lowest possible free energy system.

Current literature exists that compares the accuracy of RNA folding software when applied to viral RNAs, and cellular RNAs, suggesting there exists no difference in accuracy between the two [25]. These investigations explore viral RNAs of various lengths, accounting for PPV, sensitivity, and F_1_ scores. However, to our knowledge, very few papers exist that address the accuracy of RNA folding software when applied to viral pseudoknotted RNA transcripts alone. Moreover, the literature does not expound on the differences in accuracy regarding MEA and MFE modalities when applied to viral pseudoknotted RNA. This paper aims to condense this investigation to a dataset of 26 short pseudoknotted sequences (20-150nt), with updated versions of existing stochastic RNA prediction algorithms. This investigation will also address MEA and MFE modalities, and which of the five-folding software (**Table 1**) are more accurate when applied to this dataset.

**Table 1.** MEA and MFE folding engines applied to viral RNA structures.

| Name of the folding engine | Method of Prediction | Thermodynamic model parameters | Pseudoknots enforced | Auxiliary parameters enforced | References |
| --- | --- | --- | --- | --- | --- |
| Chiba Institute of Technology's Vsfold5: RNA Secondary Structure Prediction Server | MEA | Jacobson-Stockmayer (standard parameters) | Yes | Kuhn length, Mg<sup>2+</sup> binding, contiguous stems, minimum stem length, length of leading stem | (26) |
| Universitat Bielefeld's BiBiServ's (pKiss) | MFE | Turner model | Yes | H-type penalty, K-type penalty, maximal pseudoknot size, minimal hairpin length, lonely base pairs | (21) |
| InstitutCurie's Kinefold | MFE | Turner model | No | Co-transcriptional Fold, Simulated molecular time, Tracing and Forcing Helices | (27) |
| NUPACK 3.0 | MFE | Turner model | No | Mg<sup>2+</sup> binding, dangling end options, input of multiple interacting | (28,29) |
| Vienna RNAfold | MFE | Turner model | No | Avoiding isolated base pairs, incorporation of G–Quadruplex formation into the structure prediction algorithm, dangling end options, addition of modified base pairs | (30) |

Note: The Vienna RNAfold engine does not compute for pseudoknots and is implemented as a negative control [31]. See supplemental information [S1-S5] to view further details regarding the auxiliary parameters enforced in each of the software.

## Materials and Methods

### 2.1. RNA Folding Engines

The five RNA secondary structure prediction servers used in this paper are listed in the table below:

### 2.2. RNA Classes & Genuses Assayed

The knowledge we possess pertaining to pseudoknots and their metabolic functions holds much of its origins in the study of viral biology, mounted on well-studied strains such as flaviviruses, influenzas, and mosaic viruses [32–34]. These structures, (**Fig. 2**) encompass the regulatory elements of some viruses, controlling various phases of gene expression and function. Though many forms of pseudoknot classification have been conjectured [24,35,36] for this paper, all pseudoknots with fall under one of the following six categories (building upon the grouping proposed by Audrey Legendre et al. [24]:

1.) H-Pseudoknot
2.) HHH-Pseudoknot (kissing hairpin)
3.) HLout-Pseudoknot
4.) HLin-Pseudoknot
5.) HLout,HLin Pseudoknot
6.) LL-Pseudoknot

**Figure 2.**
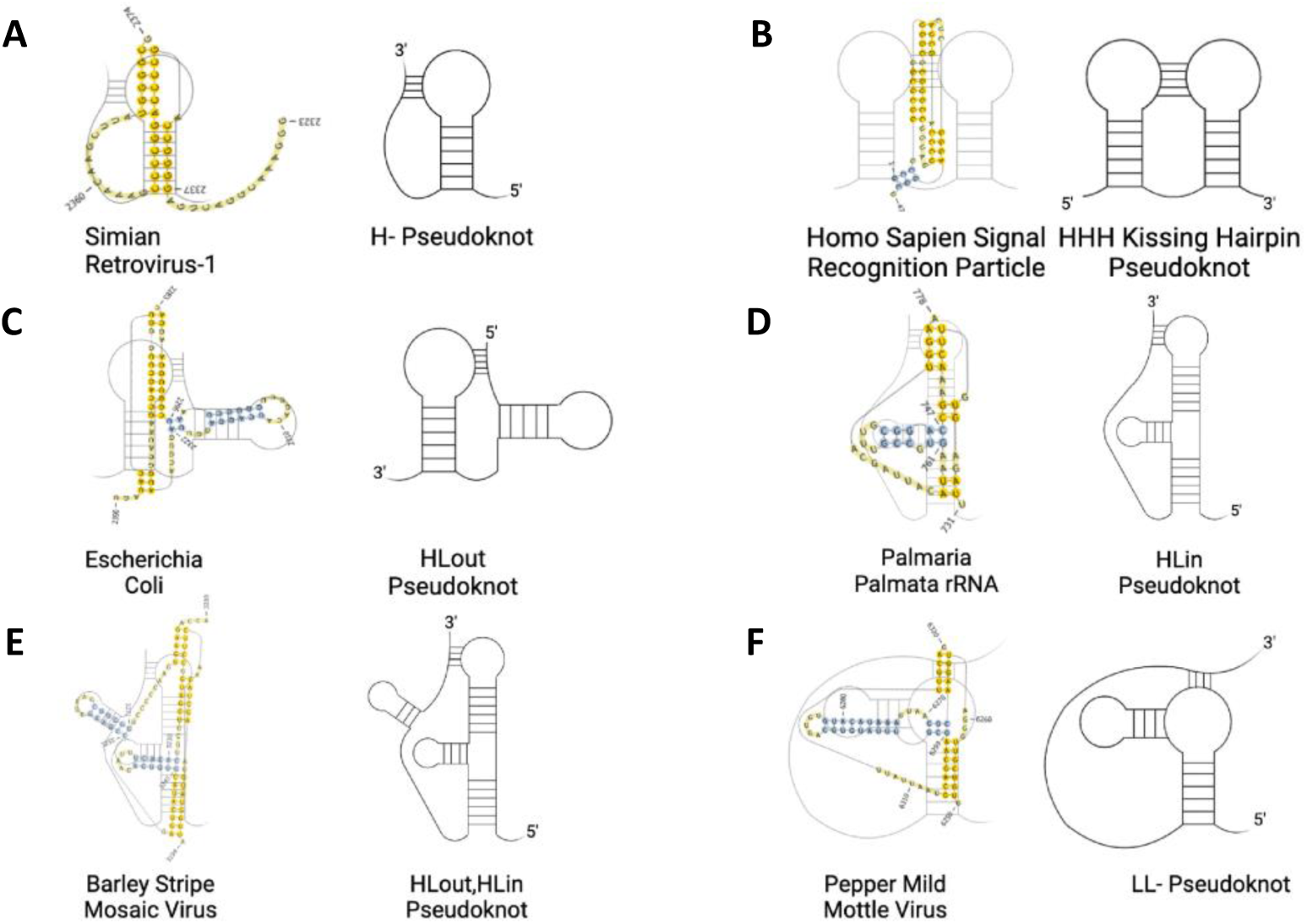
Six classes of pseudoknot. The figure depicts the class of pseudoknot (skeletal structure) on the right, with an example of an accepted structure on the left. (**A**) Simian Retrovirus-1 [37]. (**B**) Homo Sapien Signal Recognition Particle [38]. (**C**) Escherichia Coli [39]. (**D**) Palmaria Palmatra rRNA [40]. (**E**) Barley Stripe Mosaic Virus [14]. (**F**) Pepper Mild Mottle Virus [14].

Given that most classified pseudoknots in databanks, both proteins nucleic acids, are of a genus 1 [41], all pseudoknots expounded on in this investigation will have a genus of 1. The ‘genus’ of a pseudoknot, refers to a mathematical concept that correlates to the topology of a surface. The genus is a positive integer value, which corresponds with the minimum number of handles of the embedding surface of a structure, or more simply, the number of times the RNA molecule intersects with itself in three-dimensional space. A genus of 0 means that the graph can be drawn without any crossing on a sphere. A genus of 1 means that the graph can be drawn on a torus (doughnut shape), without any base pairs crossing, and so on.

### 2.3 Generating Viral Pseudoknotted RNA PseudoBase++ Dataset

Of the 398 RNAs found within the Pseudobase++ database, 205 peer-reviewed articles referenced in these sites were vetted thoroughly by the “Preferred Reporting Items for Systematic Reviews and Meta-Analyses” (PRISMA) guidelines [44]. The search identified 87 eligible studies. In addition to solely using pseudoknots of a genus 1 class, the computed MFE of all pseudoknots was also considered (see supplemental **Table S2**). Any pseudoknot resulting in too high an MFE (P<0.05) was excluded from the dataset and considered an outlier. What remained were 26 RNAs of varying sizes between 20-150nt, conforming to one of the 6 structures depicted in **Fig. 2** (see supplemental **Table S1** for further information on individual structures). Structures were derived from both plant and animal species with each structure corresponding to a unique viral genome.

As mentioned previously, all pseudoknots expounded in this report fall under at least one of the six categories described in **Fig. 2**. Of the 26 RNAs assessed, 17 consist of H-type pseudoknots, while the other nine fall under one of the other configurations listed. This skewness is intentional, and true to nature, given that hairpin-type pseudoknots (H-type) are more common by far [6]. In addition, of the pseudoknotted RNA assessed, 16 will harbor viral tRNA-like motifs, 5 will harbor viral 3’ UTR-like motifs, and 4 will harbor viral frameshifts. Each motif plays an essential role in the replication of viruses and was thus included in the dataset (see supplemental **Table S1** for further information). For example, the pseudoknotted UTRs of positive-strand RNA viruses regulate and initiate protein synthesis and replicase enzymes, L-shaped pseudoknots that resemble tRNA’s propagate viral proteins, and viral frameshifts result in the production of unique proteins (especially in retroviruses) [6].

### 2.4 Performance Metrics

Base pairings were deemed correct following David H. Matthews’s parameters [45]. Through this system, a base pair in a sequence of length N, between base pairs *i* and *j* (where 1 ≤ *i* < *j* ≤ N), would be considered correct if *i* was paired to either *j, j*-1, or *j* + 1, or if j was paired with *i, i* – 1, or *i* + 1. This model allows for some elasticity and leniency in prediction, deeming base pairing correct even if they are displaced by one nucleotide either up or downstream. This is the standard for in silico RNA computation. Further benchmarks employed by (Mathews et al. 2019) include:

- A large set of well-established reference/accepted structures to compare against experimentally derived data.
- Tests for statistical significance.
- Different RNA families/ RNA types (see supplemental **Table S1**) should be used than those used to train the methods being benchmarked.

Firstly, the percent error was used to assess the prediction accuracy of total base pairs and knotted base pairs. The percent error is expressed as the absolute value of the difference between base pairings derived from one of the five RNA folding engines (**Table 1**), and base pairs from the PseudoBase++ database derived from sequence comparison, structure probing, and NMR, put in percent format:

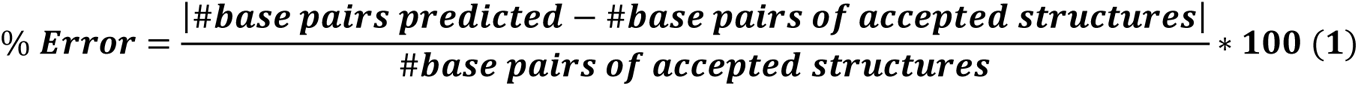

This manner of assessment is better suited to delineating between software that can or cannot predict pseudoknots. However, this metric does not encompass a software’s ability to holistically assess both knotted and total base pairs in tandem. For this reason, many papers are instead opting for sensitivity (recall) and PPV (precision) [45,46].

The sensitivity and the PPV are common, and important systems of measurement when predicting software accuracy. The former assesses a folding engine’s ability to identify correct base pairs, while the latter assesses a software’s propensity to incorrectly identify base pairs (resulting in a value of 1 to 0).

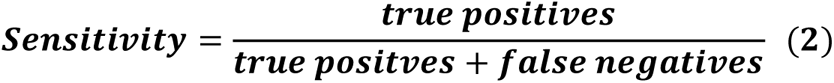

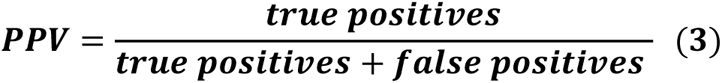

Here, sensitivity is reduced from 1 when an RNA folding engine misses pairings, while the PPV is reduced from 1 the more an RNA folding engine predicts bases that are not of the original secondary structure. Though these metrics are commonplace in almost all branches of bioinformatics, such as genomic variant calling, and drug targeting prediction [47,48] this report will impose them onto folding engines.

Once both these values have been calculated, we can derive the F_1_ scores for each structure. F_1_ scores are a standard method used to evaluate prediction software [24,45,46,49], and assess the harmonic mean between the sensitivity and the PPV by taking three of the four confusion matrix categories into account (true positive, false positive, and false negative). F_1_ scoring remains an archetype for binary classification problems [49], and integrates the boons of sensitivity and PPV to produce a higher caliber performance metric:

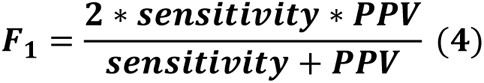

Statistical analysis including normality and lognormality tests, outlier identification, and 2-way ANOVA testing, was performed using GraphPad Prism v.9.5 software (Graph-Pad Software, San Diego, CA, USA), and accuracy metrics were reported as decimals.

## Results

The primary aims of this report are to evaluate how MEA folding modalities compare to MFE folding modalities in the context of pseudoknotted viral RNAs. While differences in the percent error of “total base pairs” predictions were non-significant across all software, the differences in the percent error of knotted base pair predictions were vastly different across the software.

The pKiss MFE folding engine exhibited the highest prediction accuracy across all MFE and MEA folding engines, and across all performance metrics used (percent error of total base pairs and knotted base pairs, sensitivity, PPV, F_1_ scores). In contrast, Kinefold exhibited the lowest values for sensitivity, PPV, and F_1_ scoring, even when compared to the control.

### 3.1 Assessment of Percent Error of in Total Base Pairs and Knotted Base Pairs

The first metric used to evaluate the accuracy of the five RNA folding engines is the percent error of total base pairs and knotted base pairs. These values were computed first by considering all total base pairs in the given model (purple), and then again by considering only pseudoknotted base pairs (blue) (**Fig. 4**). No significant difference in percent error was observed between the total number of base pairs computed by all RNA folding algorithms, with the largest discernible difference between Vsfold5 and Vienna (Adjusted P value = 0.9989). This agrees with current studies, as the challenge biotechnicians face instead lies within predictions in O(N3) and O(N4) time and space where N is the sequence length (using big O notation) [50]. While certain algorithms can reduce higher-ordered structures to O(N3), thereby reducing computational complexity, a growing minimal N value correlates with more possible pseudoknots, making the algorithm less accurate [21].

**Figure 3.**
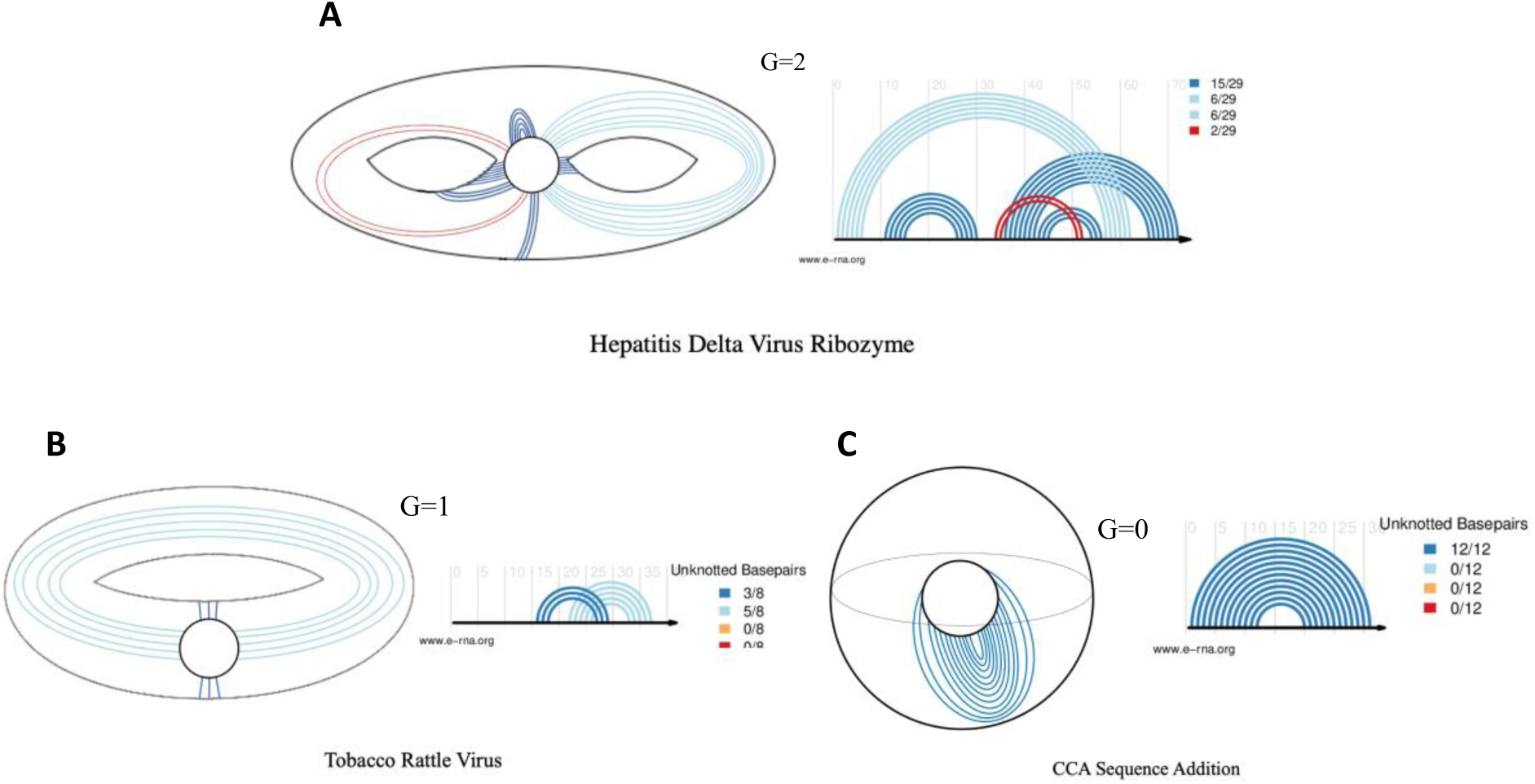
Depiction of genus 0,1, and 2 RNA’s. (**A**) Hepatitis delta virus ribozyme in 3D space, and a planer arc and chord diagram [41,42]. (**B**) Tobacco rattle virus in 3D space, and a planer arc and chord diagram [14]. (**C**) CCA sequence adding polymerase in 3D space, and a planer arc and chord diagram [43]. The search identified 87 eligible studies. In addition to solely using pseudoknots of a genus 1 class, the computed MFE of all pseudoknots was also considered (see supplemental **Table S2**). Any pseudoknot resulting in too high an MFE (P<0.05) was excluded from the dataset and considered an outlier. What remained were 26 RNAs of varying sizes between 20-150nt, conforming to one of the 6 structures depicted in Fig. 2 (see supplemental **Table S1** for further information on individual structures). Structures were derived from both plant and animal species with each structure corresponding to a unique viral genome.

**Figure 4.**
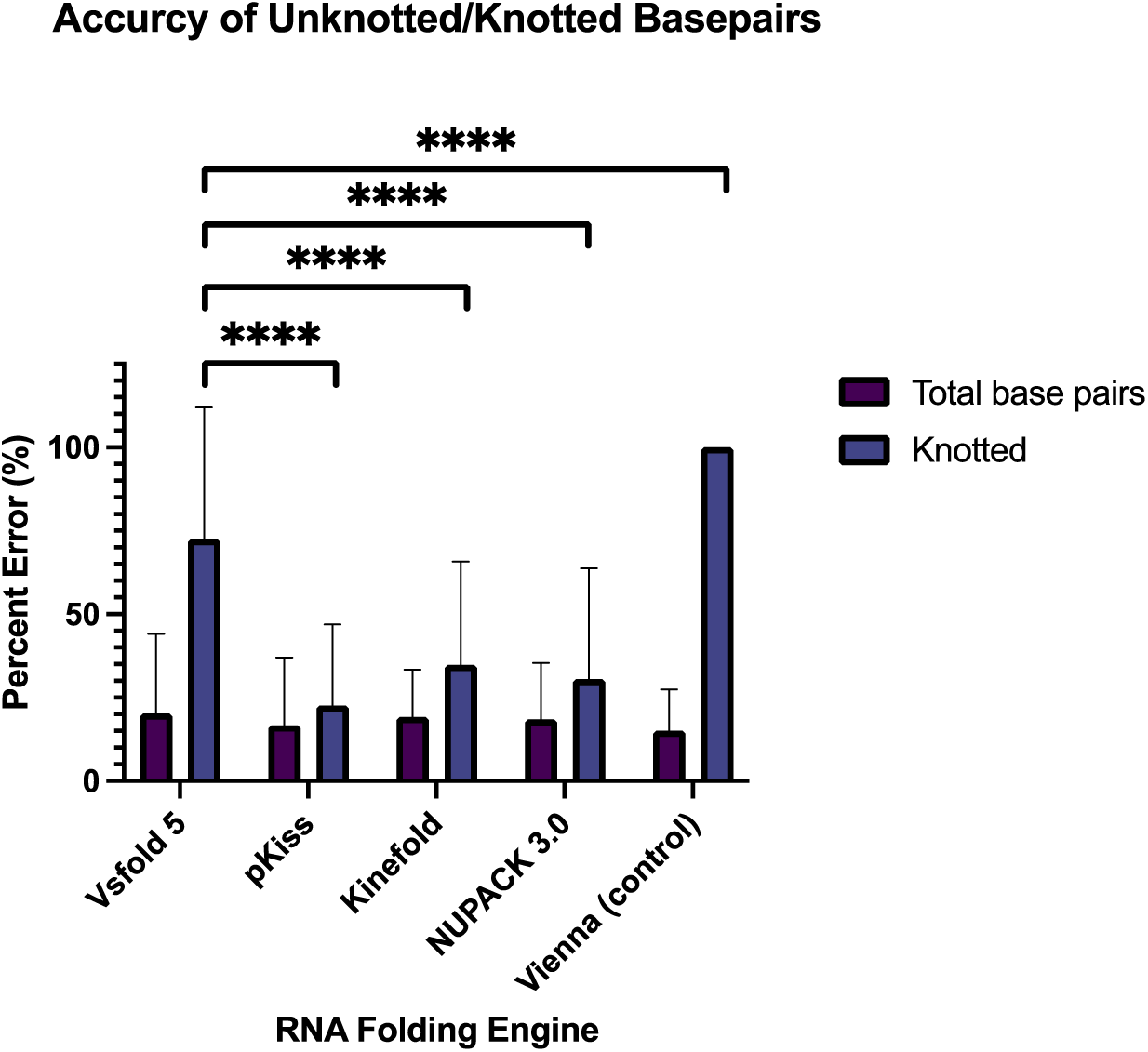
Percent error of total base pairs and knotted base pairs. Data corresponds to the mean percent error ± SD. Mean percent error ± SD, across all four MFE RNA folding engines, was compared to that of the MEA computations of Vsfold 5, with statistical analysis performed using a two-way ANOVA, followed by a Tukey’s multiple comparisons test. **** (P ≤ 0.0001). ROUT test was instated to identify outliers (Q=1%).

The mean (x̄ ) of the percent error of total base pairs generated by each software was 17.95%, with Vsfold 5 exhibiting the highest (x̄ =20.23 ± 23.94%), and the negative control exhibiting the lowest (x̄ =15.07 ± 12.35%), which was expected. Though current literature now advocates for software that computes MEA, rather than MFE [24], the Vienna package is specifically designed to predict planer secondary structures.

A much higher percent error was exhibited in knotted viral RNA structures produced by the MEA engine Vsfold 5, relative to its MFE computing counterparts (apart from the negative control). Vsfold 5, was expected to give the lowest percent error. However, the lowest percent error for knotted base pairs was instead exhibited by MFE structures generated by pKiss (x̄ =22.37 ± 24.2%), while Vsfold 5 retained a mean percent error of (69.91 ±39.30%). While the low percent error exhibited by pKiss could be the result of the pseudoknot “enforce” constraint, it is more likely that this outcome was multivariable, equating to the Turner energy model used, and the sensitive auxiliary parameters enforced by the program (refer to **Table 1** and supplementary information **S2**).

### 3.2 Sensitivity and Positive Predictive Value (PPV) of Folding Engines

The second metric used to assess RNA folding engines was sensitivity, coupled with positive predictive value (PPV). With all 26 viral RNAs considered, the highest mean sensitivity and PPV were derived from pKiss, with values of (x̄ =0.88 ± 0.14) and (x̄ =0.82 ± 0.16) respectively. These values trump those of prior versions, reporting lesser sensitivity (x̄ =0.80 ± 0.24), and PPV values (x̄ =0.75 ± 0.27) [24]. Conversely, the lowest mean sensi-tivity and PPV were derived from Kinefold, with values of (x̄ =0.14 ± 0.23) and (x̄ =0.171 ± 0.31) respectively.

It is important to emphasize the increase in both PPV and sensitivity of the newer folding engines listed in (**Table 1**) when compared to older folding engines previously reported in the literature. Such examples may include ProbKnot, with a sensitivity of x̄ =0.693 and a PPV of x̄ =0.613 [51], and PKNOTS (the older version of pKiss), with older papers reporting (sensitivity: x̄ =0.828, PPV: x̄ =0.789), and newer papers reporting (sensitivity: x̄ =0.855, PPV: x̄ =0.808) [52]. Papers promoting RNA software that shares MEA, and MFE properties, such as BiokoP dating back five years prior have also reported a lesser sensitivity (0.81 ± 0.22) and PPV values (0.75 ± 0.26) [24]. Note that confounding variables between this paper and the referenced literature are minimal, as all reports screened for a diverse set of RNA pseudoknots. Ultimately, pKiss outperformed the mean of the Vsfold 5 MEA prediction software by x̄ =0.296.

We would like to note that PPV values of free energy minimization, that is, MFE folding engines, have been shown to be lower than sensitivity values [53,54]. This is likely because structures accepted in the literature can be missing base pairs that may occur experimentally, and because the thermodynamics imposed by MFE algorithms often overshoot the number of canonical base pairings (because it is the formation of base pairs that innately lowers the Gibbs free energy of a structure) [55]. However, this trend does not present itself in all four experimental conditions, only in pKiss and NUPACK 3.0. This is because updated software such as this implements has more accurate assessment of the thermodynamic properties of the structure, removing unwanted pairs and improving overall performance [51].

### 3.3 Quality of Prediction Software Assessed Via F_1_ scoring

F_1_ scores were derived from the sensitivity and PPV values (**Fig. 6**). This modality is often adapted to assess prediction accuracy, so long as the reference structure is given. Of all average F_1_ scores generated from all five stochastic folding algorithms, the pKiss engine computed the largest values (x̄ = 0.844 ± 0.138), while Kinefold computed the lowest values (x̄ =0.150 ± 0.273). Of all MFE folding algorithms used, pKiss was the only one to significantly outperform the mean F_1_ score of the Vsfold 5 MEA engine by a value of 0.235. These values exceed those of previously reported folding engines, such as CCJ and ProbKnot, with F_1_ scores of (x̄ =0.644) and (x̄ =0.738) respectively, while at the same time, underperforming when compared to deep learning algorithms like ATTfold with an F_1_ score of (0.966) [46,56]. Outliers were found in Vsfold 5, Kinefold, and NUPACK 3.0 packages, corresponding to large data sets.

**Figure 5.**
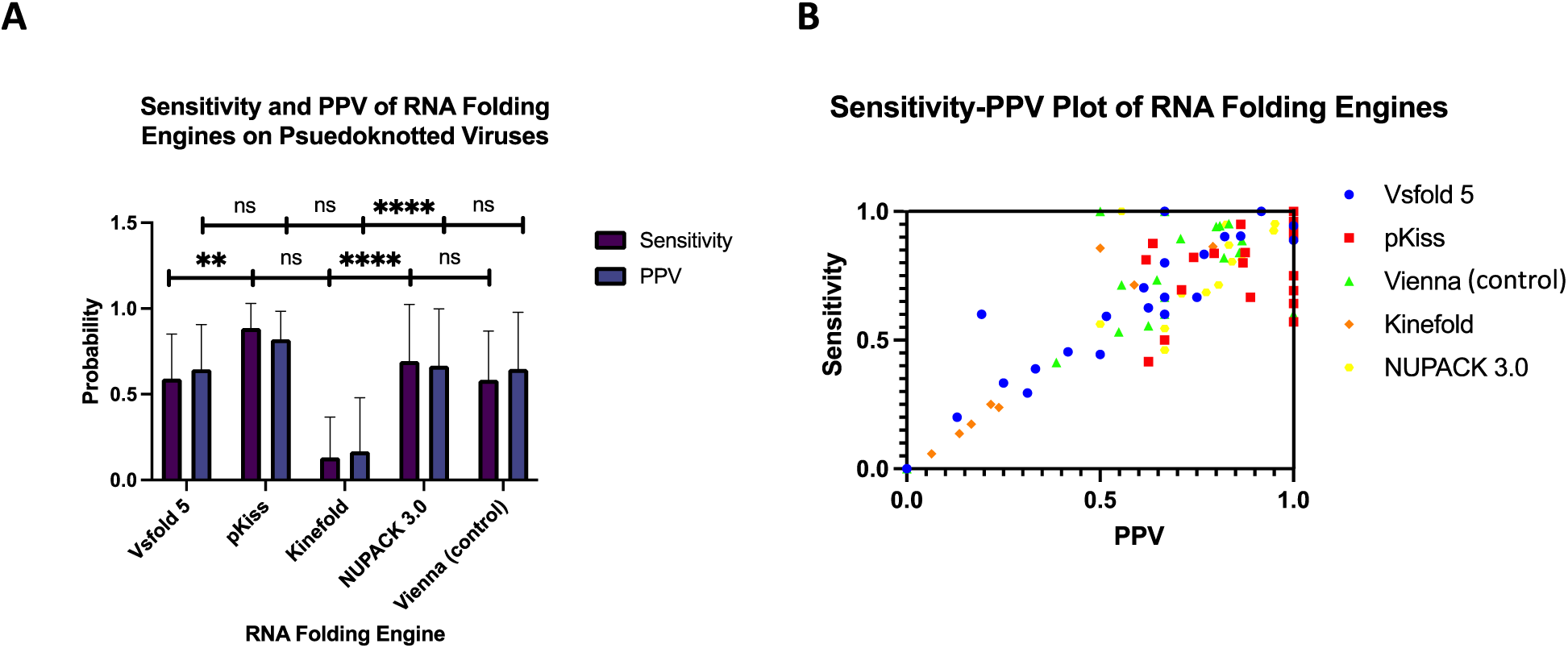
Sensitivity and PPV of RNA folding engines on pseudoknotted viruses. (**A**) Data corresponding to the mean sensitivity and PPV ± SD across all five experimental conditions were compared to the sensitivity and PPV generated by Vsfold 5. Statistical analysis was performed using a two-way ANOVA, followed by a Tukey’s multiple comparisons test. **** (P ≤ 0.0001), ** (P ≤ 0.0021), and ns (P ≥ 0.1234). ROUT test was instated to identify outliers (Q=1%). (**B**) Sensitivity and PPV values plotted in a 1x1 matrix.

**Figure 6.**
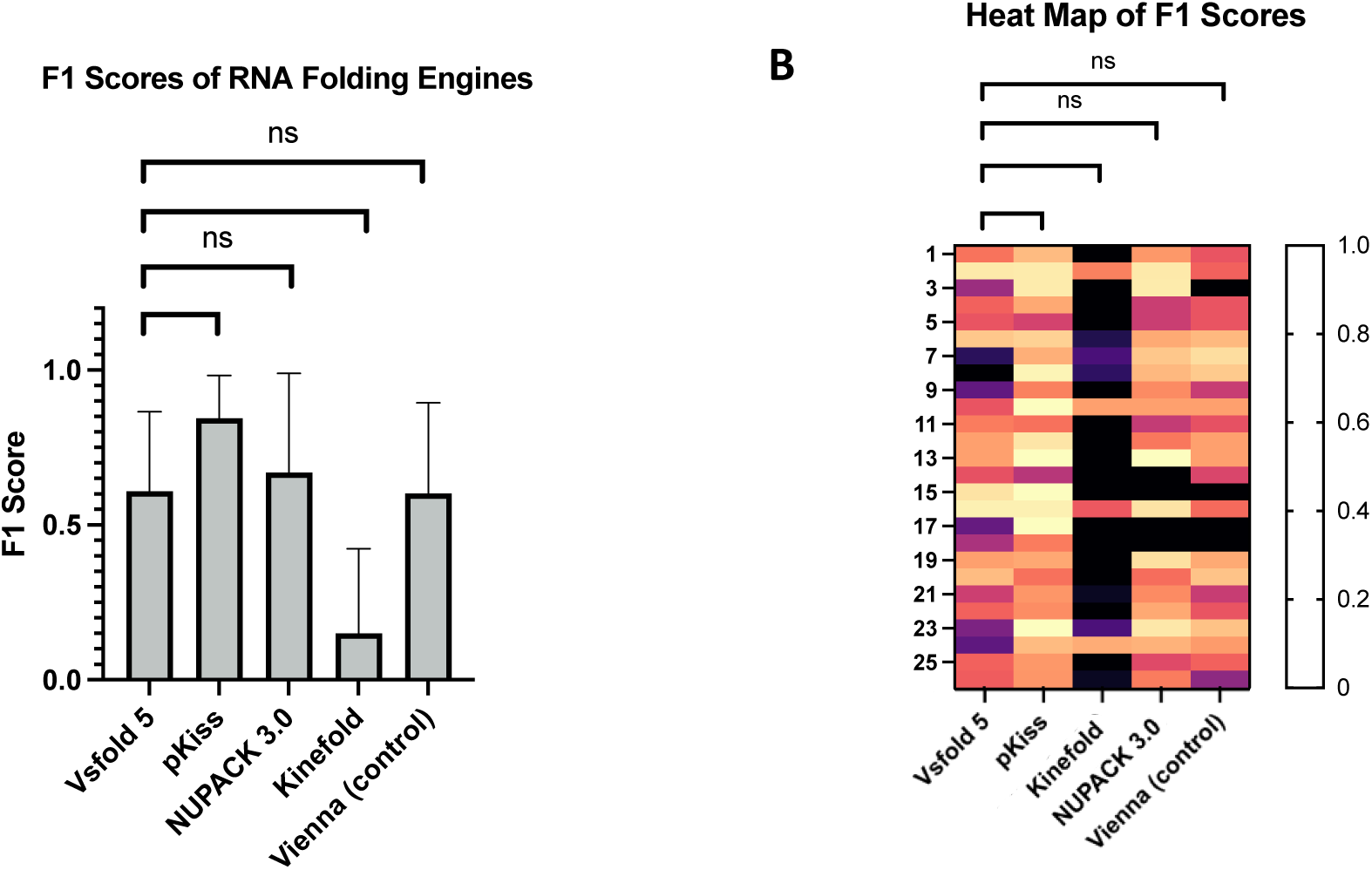
F_1_ scores of virus database. (**A**) Mean F_1_ Scores ± SD depicted as a bar graph. (B) Individual F_1_ scores generated by all software are depicted as a heatmap, with white equating to the best performance F1 =1, and black equating to the worst performance F_1_ = 0. Data corresponding to the mean F_1_ scores ± SD. Mean F1 scores ± SD, across all five experimental conditions, were compared to that of the F_1_ scores of Vsfold 5, with statistical analysis performed using a two-way ANOVA, followed by a Tukey’s multiple comparisons test. **** (P ≤ 0.0001), ** (P ≤ 0.0021), and ns (P ≥ 0.1234). ROUT test was instated to identify outliers (Q=1%).

## Discussion

The underlying functions and computational modalities for RNA prediction algorithms have greatly evolved since Nussinov’s dynamic programming algorithm, [57] a formalism derived in 1978, and arguably the genesis of predictive RNA folding whereby:

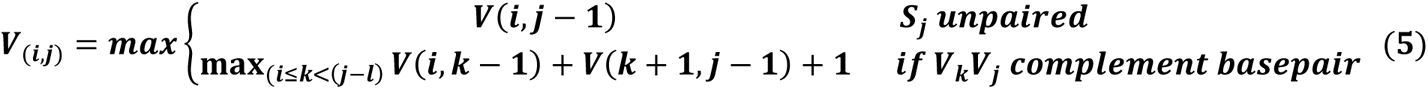

Where:

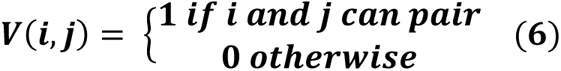

Using these systems, it is possible to generate minimum free energy (MFE) predictions in kcal/mol of a structure via tracebacking, resulting in a probabilistic model of the nature of the RNA template in vivo. This original scheme has been amended many times over, with Zuker’s algorithm (1981) being the most notable change-an amendment that is still used to this day [58].

The realization of these formalisms coincides with the discovery of the first pseudoknotted plant viruses discovered in the early 1980’s. Today, they make up many of the pseudoknots found in various online databases/databanks and are recognized as common motifs that allow for viral mRNA function, ribosome function, and replication [6]. The inherent vastness of viral RNA in nature, and consequently, within pseudoknot databases (like those taken from the PseudoBase++ [23] and RNA STRAND [59]), is the main reason for this investigation. We have provided evidence suggesting that MEA software is not always the optimal method of topological prediction when applied to short viral pseudoknotted RNA. Moreover, minor improvements have been made to previous MFE reports, while underperforming existing deep learning and machine learning algorithms.

Of the metrics that were employed under the remits of this investigation, none demonstrated that the MEA algorithm used, Vsfold 5, was inherently superlative to its MFE counterparts. This suggests that some short viral pseudoknotted RNA’s (20-150nt) may often result in their lowest free energy model (granted that salinity, Mg^2+^ conc., and other environmental variables remain constant). This conclusion is shared amongst pseudoknotted viral RNAs that are of different structures (e.g. H-type, LL-type), and of different motifs (e.g. Viral tRNA-like, Viral 3 UTR).

It should be noted that the thermodynamics within the cell, as well as the many auxiliary folding pathways of RNA, become muddled when exploring the extensive cellular environment in vivo. This is where *ab initio* and comparative approaches come into play. However, one should consider that *in silico* studies will oftentimes lack direct cellular relevance, as researchers remain aloof to the broader physiological consequences of change [5]. Therefore, it should be emphasized that both in vitro, and *in silico* approaches are necessary to explore the nature of viral RNA’s.

## Conclusions

To our knowledge, this paper embodies the first attempt at applying a suite of RNA folding engines to a dataset solely comprised of viral pseudoknotted RNAs. The data computed in this paper is founded upon different MEA and MFE software that have received updates in recent years, and the accuracy of these RNA folding engines was benchmarked following David H. Mathews’ parameters. The evidence provided suggests that viral pseudoknotted RNAs may conform to the MFE structure in some cases, rather than the MEA structure. Under the scope of these quality folding engines, pKiss provided the most accurate structures when compared to data experimentally derived from mutagenesis, sequence comparison, structure probing, and NMR, while Kinefold resulted in the least accurate structures. This infers that the veracity of the underlying thermodynamic model parameters (e.g. Turner model, Jacobson-Stockmayer), is compromised if the auxiliary parameters are not enforced (e.g. Mg^2+^ binding, dangling end options, H-type penalties).

To expedite the screening of RNAs, whether they be knotted or planer, we must achieve a better understanding of the thermodynamics associated with cellular processes, and how they govern the shaping of RNA. The explored *ab initio* methodologies provide more accurate results than previously reported, though they do not outperform deep learning algorithms. The exploration of RNA outside the wet lab might seem counterintuitive, the computing power we now possess lends to efficacious predictions. Limitations are present in both in vitro, and *in silico* methodologies. However, both are necessary to further the exploration of drug targets, mRNA vaccines, thermosensors, and RNA-based genome editing.

## Supplementary Materials

The data referenced in supplementary materials can be found at _______.

## Author Contributions

Conceptualization, V.M., methodology, V.M., formal analysis. V.M., M.C., data curation, V.M., M.C., writing-original draft preparation, V.M., review and editing, V.M., S.Z., E.E., J.P. All authors have read and agreed to the published version of the manuscript.

## Informed Consent Statement

Not applicable.

## Data Availability Statement

The data in this study is present within the present article, as well as the supplementary materials.

## Acknowledgments

We would like to thank Ajay Singh, Jamie M. Robertson, and the Effective Writing for Healthcare program (Harvard Medical School), for reviewing and editing this work. We thank the Das and Barna laboratories (Stanford University), as well as the entirety of the Eterna community for their contributions to the project. We would like to thank Henri Orland for providing supplemental aid to the content of the paper. Finally, we would like to acknowledge the Eterna OpenKnot labs project for inspiring this work, and the National Instate of Health for generating their Pseudokbase++ database.

